# *pals-22*, a member of an expanded *C. elegans* gene family, controls silencing of repetitive DNA

**DOI:** 10.1101/145094

**Authors:** Eduardo Leyva-Díaz, Nikolaos Stefanakis, Inés Carrera, Lori Glenwinkel, Guoqiang Wang, Monica Driscoll, Oliver Hobert

## Abstract

Repetitive DNA sequences are subject to gene silencing in various animal species. Under specific circumstances repetitive DNA sequences can escape such silencing. For example, when exogenously added, extrachromosomal DNA sequences that are stably inherited in multicopy repetitive arrays in the nematode *C. elegans* are frequently silenced in the germline, whereas such silencing often does not occur in the soma. This indicates that somatic cells might utilize factors that prevent repetitive DNA silencing. Indeed, such “anti-silencing” factors have been revealed through genetic screens that identified mutant loci in which repetitive transgenic arrays are aberrantly silenced in the soma. We describe here a novel locus, *pals-22* (for <u>p</u>rotein containing <u>ALS</u>2CR12 domain), required to prevent silencing of repetitive transgenes in neurons and other somatic tissue types. *pals-22* deficiency also severely impacts animal vigor and confers phenotypes reminiscent of accelerated aging. We find that *pals-22* is a member of a large family of divergent genes (39 members), defined by the presence of an ALS2CR12 domain. While gene family members are highly divergent, they show striking patterns of genomic clustering. The family expansion appears *C. elegans-*specific and has not occurred to the same extent in other nematode species. Previous transcriptome analysis has revealed that most of the *pals* genes are induced under stress conditions or upon infection by intracellular parasites. The transgene silencing phenotype observed upon loss of cytoplasmically localized PALS-22 protein depends on the biogenesis of small RNAs, since silencing is abolished in the RNAi defective mutant *rde-4*, suggesting that *pals-22* might regulate RNAi dependent silencing in the cytoplasm of neurons and other tissues. We speculate that the *pals* gene family may be part of a species-specific cellular defense mechanism.

## INTRODUCTION

Over half the human genome consists of repetitive DNA elements (Lander *et al.* 2001; de Koning *et al.* 2011). The view of the role of repetitive DNA has evolved in the last decades from considering it as “junk DNA” to the recognition of repetitive DNA as essential for genome function (Doolittle AND Sapienza 1980; Orgel AND Crick 1980; Lynch AND Conery 2003; Shapiro AND von Sternberg 2005). The main constituents of these repetitive DNA elements are retrotransposons, a large family of transposable elements capable of copying themselves and reinserting into the host genome (Kazazian 2004; Goodier AND Kazazian 2008; Cordaux AND Batzer 2009). Retrotransposons and other elements with the ability to copy themselves pose a threat to genome integrity due to the potential deleterious effects of landing in coding or regulatory regions (Friedli AND Trono 2015). The activation of proto-oncogenes in some leukemias represent an example of such harmful consequences (Hacein-Bey-Abina *et al.* 2003). However, repetitive DNA elements have also been found to play beneficial roles in a number of processes, ranging from the regulation of gene expression to interaction with nuclear structures for genome packaging, to DNA repair and restructuring (Shapiro AND von Sternberg 2005; Goke AND Ng 2016). Moreover, retrotransposons are a source of regulatory elements and alternative gene isoforms. Hence, transposable elements have been postulated as a powerful genetic force involved in the evolution of organismal complexity (Kazazian 2004; Friedli AND Trono 2015).

To balance the deleterious and beneficial features of repetitive sequences, organisms have evolved ways to finely tune the regulation of repetitive DNA elements (Schlesinger AND Goff 2015; Chuong *et al.* 2017). For example, endogenous silencing mechanisms have evolved to prevent genome damage by the spread of mobile repetitive DNA elements. Silencing is achieved by DNA binding proteins, histone modifications and DNA methylation, both of which can be directed by sequence-specific repressive transcription factors and by small RNAs. For example, retrotransposons are extensively recognized by the Krueppel-associated box-zinc finger (KRAB-ZFP) proteins (Rowe *et al.* 2010; Quenneville *et al.* 2012; Jacobs *et al.* 2014), which form a large family of repressive transcription factors. These transcription factors are among the fastest evolving group of genes in the human genome and their diversity facilitates their ability to recognize a large number of retrotransposons (Nowick *et al.* 2010). Sequence-specific binding to retrotransposons by KRAB-ZFP factors triggers a cascade leading to chromatin-based silencing mechanisms (Wolf AND Goff 2009).

RNAs of retrotransposons that escape transcriptional silencing are targeted and destroyed by the small RNA pathways in the cytoplasm (Toth *et al.* 2016). RNA-based mechanisms represent the most ancient defense against the genomic spread of repetitive DNA elements (Friedli AND Trono 2015). These mechanisms comprise the action of small RNA molecules, including small interfering RNAs (siRNAs), PIWI-interacting RNAs (piRNAs) and microRNAs (miRNAs), which guide repressor protein complexes to particular targets in a sequence-specific manner. In addition to intervening at the post-transcriptional level, small RNAs can also intervene at the transcriptional level by directing deposition of repressive histone marks and DNA methylation to copies of retrotransposons and other elements (Le Thomas *et al.* 2013). piRNAs and siRNAs can be produced from repetitive DNA elements, which they silence in return (Law AND Jacobsen 2010). Thus, in many organisms, repetitive DNA serves as a trigger for gene silencing.

Repetitive DNA elements are not only abundant in vertebrate genomes. Repetitive DNA elements also abound in model organisms with comparatively smaller genome sizes: repetitive DNA accounts for 34-57% of the total genome in *Drosophila melanogaster* (Celniker *et al.* 2002), and at least 17% of the *Caenorhabditis elegans* genome (Stein *et al.* 2003). Repetitive DNA elements can also be generated experimentally. DNA transformation techniques in *C. elegans* produce repetitive extrachromosomal DNA arrays (“simple” arrays)(Mello *et al.* 1991). Several studies have shown that expression of transgenes organized in these repetitive arrays is silenced both in somatic cells and in the germline through heterochromatin formation, involving several chromatin factors (reviewed in (Cui AND Han 2007)). Somatic, and especially, germline transgene expression can be improved when the transgenic DNAs are cotransformed with an excess of carrier DNA, producing a less repetitive, more “complex” array (Kelly *et al.* 1997).

Importantly, gene expression from repetitive genomic regions can still be observed, suggesting there are mechanisms that can modulate silencing effects (Tseng *et al.* 2007). Multiple forward genetic screens in *C. elegans* have indeed identified factors that act to counter silencing of genes contained in repetitive sequences, based on screens for mutations that alter the activity of transgenes present in tandemly repeated arrays. (Hsieh *et al.* 1999; Grishok *et al.* 2005; Tseng *et al.* 2007; Fischer *et al.* 2013). In a classic study, mutations in *tam-1* (a RING finger/B-box factor) were found to reduce the expression of transgenes organized in simple but not complex repetitive arrays (Hsieh *et al.* 1999). Therefore, *tam-1* is an “anti-silencing factor” that attenuates the context-dependent silencing mechanism affecting multi-copy tandem-array transgenes in *C. elegans.* A subsequent study identified mutations in another gene important for expression of repetitive sequences, *lex-1*, which genetically interacts with *tam-1* (Tseng *et al.* 2007). LEX-1 encodes a protein containing an ATPase domain and a bromodomain, both of which suggest that LEX-1 associates with acetylated histones and modulates chromatin structure. Hence, TAM-1 and LEX-1 are anti-silencing factors that function together to influence chromatin structure and to promote expression from repetitive sequences. Further studies found that tandem-array transgenes become silenced in most mutants that cause enhanced exogenous RNAi (Simmer *et al.* 2002; Kennedy *et al.* 2004; Fischer *et al.* 2013). Examples of gene inactivations known to cause increased transgene silencing and enhanced RNAi include the retinoblastoma like gene *lin-35* (Hsieh *et al.* 1999; Wang *et al.* 2005; Lehner *et al.* 2006), the RNA-dependent RNA polymerase *rrf-3* (Simmer *et al.* 2002), and the helicase gene *eri-6/7* (Fischer *et al.* 2008). Silencing of repetitive DNA elements (multi-copy transgenes) depends on a complex interaction between different small RNA pathways (Fischer *et al.* 2013).

Here, we identify a novel locus, *pals-22*, whose loss confers a transgene silencing phenotype. *pals-22* mutants display context-dependent array silencing, affecting the expression of highly repetitive transgenes but not single copy reporters. Animals lacking *pals-22* show locomotory defects and premature aging. *pals-22* is a member of a large family of divergent genes defined by the presence of an ALS2CR12 domain. The ALS2CR12 domain protein family is specifically expanded in *C. elegans*, and *pals* gene family members are clustered in the genome. We found that transgene silencing on *pals-22* mutants depends on the RNAi pathway, indicating that *pals-22* might act as regulator of small RNA-dependent gene silencing.

## MATERIALS AND METHODS

### Mutant strains

Strains were maintained by standard methods (Brenner 1974). The *C. elegans* mutant alleles used in this study were: *pals-22(ot723), pals-22(ot810), pals-22(ot811), rde-4(ne301)* (Tabara *et al.* 1999) and *tam-1(cc567)* (Hsieh *et al.* 1999).

### Reporter and transgenic strains

The *C. elegans* transgenic strains used in this study were *otIs381[ric-19^prom6^::NLS::gfp]*, *otIs380[ric-19^prom6^::NLS::gfp]*, *ccls4251[myo-3^prom^::gfp]*, *otIs251[cat-2^prom^::gfp]*, *otIs355[rab-3^prom1^::NLS::rfp]*, *otIs447[unc-3^prom^::mChOpti]*, *otEx6944 [ric-4^prom26^::NLS::yfp]*, *otIs620[unc-11^prom8^::NLS::gfp]*, *otIs353[ric-4^fosmid^::SL2::NLS-YFP-H2B]*, *otIs534[cho-1^fosmid^::SL2::NLS-YFP-H2B]*, *otTi32[lin-4^prom^::yfp]*, *ieSi60[myo-2^prom^::TIR1::mRuby]*, *otEx7036[pals-22^prom^::gfp]*, *otEx7037 [pals-22::gfp]. pals-22* GFP reporters were generated using a PCR fusion approach (Hobert 2002). Genomic fragments were fused to the GFP coding sequence, which was followed by the *unc-54* 3′ UTR. See Table 1 for transgenic strain names and microinjection details.

**Table 1:**
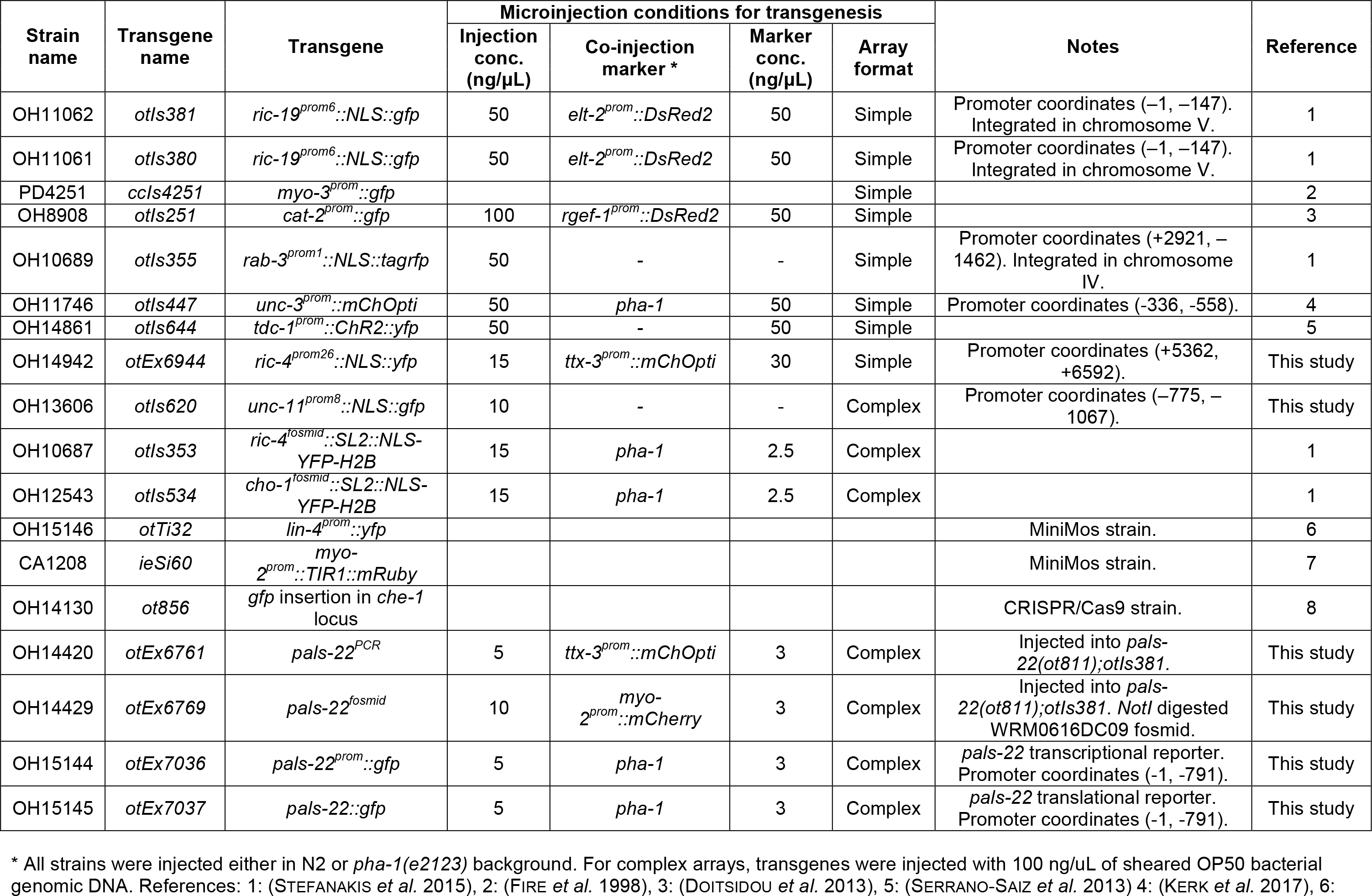
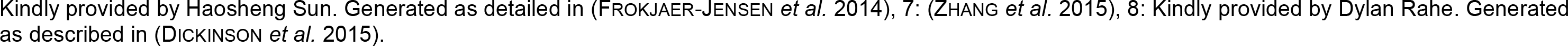
Transgenic strains used in this study.

| Strain name | Transgene name | Transgene | Microinjection conditions for transgenesis |  |  |  | Notes | Reference |
| --- | --- | --- | --- | --- | --- | --- | --- | --- |
|  |  |  | Injection conc. (ng/μL) | Co-injection marker * | Marker conc. (ng/μL) | Array format |  |  |
| OH11062 | *otIs381* | *ric-19<sup>prom6</sup>::NLS::gfp* | 50 | *elt-2<sup>prom</sup>::DsRed2* | 50 | Simple | Promoter coordinates (–1, –147). Integrated in chromosome V. | 1 |
| OH11061 | *otIs380* | *ric-19<sup>prom6</sup>::NLS::gfp* | 50 | *elt-2<sup>prom</sup>::DsRed2* | 50 | Simple | Promoter coordinates (–1, –147). Integrated in chromosome V. | 1 |
| PD4251 | *ccls4251* | *myo-3<sup>prom</sup>::gfp* |  |  |  | Simple |  | 2 |
| OH8908 | *otIs251* | *cat-2<sup>prom</sup>::gfp* | 100 | *rgef-1<sup>prom</sup>::DsRed2* | 50 | Simple |  | 3 |
| OH10689 | *otIs355* | *rab-3<sup>prom1</sup>::NLS::tagrfp* | 50 | - | - | Simple | Promoter coordinates (+2921, –1462). Integrated in chromosome IV. | 1 |
| OH11746 | *otIs447* | *unc-3<sup>prom</sup>::mChOpti* | 50 | *pha-1* | 50 | Simple | Promoter coordinates (-336, -558). | 4 |
| OH14861 | *otIs644* | *tdc-1<sup>prom</sup>::ChR2::yfp* | 50 | - | 50 | Simple |  | 5 |
| OH14942 | *otEx6944* | *ric-4<sup>prom26</sup>::NLS::yfp* | 15 | *ttx-3<sup>prom</sup>::mChOpti* | 30 | Simple | Promoter coordinates (+5362, +6592). | This study |
| OH13606 | *otIs620* | *unc-11<sup>prom8</sup>::NLS::gfp* | 10 | - | - | Complex | Promoter coordinates (–775, –1067). | This study |
| OH10687 | *otIs353* | *ric-4<sup>fosmid</sup>::SL2::NLS-YFP-H2B* | 15 | *pha-1* | 2.5 | Complex |  | 1 |
| OH12543 | *otIs534* | *cho-1<sup>fosmid</sup>::SL2::NLS-YFP-H2B* | 15 | *pha-1* | 2.5 | Complex |  | 1 |
| OH15146 | *otTi32* | *lin-4<sup>prom</sup>::yfp* |  |  |  |  | MiniMos strain. | 6 |
| CA1208 | *ieSi60* | *myo-2<sup>prom</sup>::TIR1::mRuby* |  |  |  |  | MiniMos strain. | 7 |
| OH14130 | *ot856* | *gfp* insertion in *che-1* locus |  |  |  |  | CRISPR/Cas9 strain. | 8 |
| OH14420 | *otEx6761* | *pals-22<sup>PCR</sup>* | 5 | *ttx-3<sup>prom</sup>::mChOpti* | 3 | Complex | Injected into *pals-22(ot811);otIs381*. | This study |
| OH14429 | *otEx6769* | *pals-22<sup>fosmid</sup>* | 10 | *myo-2<sup>prom</sup>::mCherry* | 3 | Complex | Injected into *pals-22(ot811);otIs381*. *NotI* digested WRM0616DC09 fosmid. | This study |
| OH15144 | *otEx7036* | *pals-22<sup>prom</sup>::gfp* | 5 | *pha-1* | 3 | Complex | *pals-22* transcriptional reporter. Promoter coordinates (-1, -791). | This study |
| OH15145 | *otEx7037* | *pals-22::gfp* | 5 | *pha-1* | 3 | Complex | *pals-22* translational reporter. Promoter coordinates (-1, -791). | This study |

\* All strains were injected either in N2 or *pha-1(e2123)* background. For complex arrays, transgenes were injected with 100 ng/μL of sheared OP50 bacterial genomic DNA. References: 1: (STEFANAKIS *et al.* 2015), 2: (FIRE *et al.* 1998), 3: (DOITSIDOU *et al.* 2013), 5: (SERRANO-SAZ *et al.* 2013) 4: (KERK *et al.* 2017), 6:
Kindly provided by Haosheng Sun. Generated as detailed in (FROKJAER-JENSEN *et al.* 2014), 7: (ZHANG *et al.* 2015), 8: Kindly provided by Dylan Rahe. Generated as described in (DICKINSON *et al.* 2015).

### Forward genetic screens

Standard ethyl methanesulfonate (EMS) mutagenesis was performed on the fluorescent transgenic reporter strain: *otIs381[ric-19^prom6^::NLS::gfp]* and ~60,000 haploid genomes were screened for expression defects with an automated screening procedure (Doitsidou *et al.* 2008) using the Union Biometrica COPAS FP-250 system. To identify the causal genes of the mutants obtained, we performed Hawaiian single nucleotide polymorphism (SNP) mapping and whole-genome sequencing (Doitsidou *et al.* 2010) followed by data analysis using the CloudMap pipeline (Minevich *et al.* 2012). The following mutant alleles were identified: *pals-22(ot810)* and *pals-22(ot811).* The mutant allele *pals-22(ot723)* was identified in an independent manual clonal screen for changes in reporter expression in neurons of *otIs381[ric-19^prom6^::NLS::gfp]* after EMS mutagenesis.

### Microscopy

Worms were anesthetized using 100 mM sodium azide (NaN_3_) and mounted on 5% agarose pads on glass slides. All images (except Figure 3, B-C, and Figure S1C) were acquired as Z-stacks of ~1 μm-thick slices with the Micro-Manager software (Edelstein *et al.* 2010) using the Zeiss Axio Imager.Z1 automated fluorescence microscope. Images were reconstructed via maximum intensity Z-projection of 2-10 μm Z-stacks using the ImageJ software (Schneider *et al.* 2012). Images shown in Figure 3, B-C, and Figure S1C were acquired using a Zeiss confocal microscope (LSM880). Several z-stack images (each ~ 0.4 µm thick) were acquired with the ZEN software. Representative images are shown following orthogonal projection of 2-10 µm z-stacks.

### Single molecule FISH

smFISH was done as previously described (Ji AND van Oudenaarden 2012). Samples were incubated overnight at 37°C during the hybridization step. The *ric-19* and *gfp* probes were designed using the Stellaris RNA FISH probe designer and was obtained conjugated to Quasar 670 from Biosearch Technologies.

### Fluorescence quantification

Synchronized day 1 adult worms were grown on NGM plates seeded with OP50 and incubated at 20°C. The COPAS FP-250 system (Union Biometrica) was used to measure the fluorescence of 200-1000 worms for each strain.

### Bioinformatic analysis

The ALS2CR12 domain phylogenetic tree was generated using MrBayes (Huelsenbeck AND Ronquist 2001; Ronquist AND Huelsenbeck 2003), lset nst=6 rates=invgamma, ngen increased until the standard deviation of split frequencies < 0.05. Input protein coding sequences for ALS2CR12 domain and PALS family orthologous proteins were aligned with M-Coffee (Wallace *et al.* 2006). The MrBayes tree figure was rendered with FigTree (http://tree.bio.ed.ac.uk/software/figtree/).

### RNAi by feeding

RNAi was performed as previously described with minor adaptations (Kamath AND Ahringer 2003). L4-stage hermaphrodite worms were placed onto NGM plates containing seeded bacteria expressing dsRNA for each assayed gene. After 24 h at 20°C, adults were removed. After a further 36-40 h at 20°C, phenotypes were scored blindly.

### Computer swim analysis

The swimming assay was performed using the CeleST program as previously described (Restif *et al.* 2014). In brief, we transferred five (day-1) adult hermaphrodites into 50 μl M9 buffer located in a 10 mm staggered ring on a glass slide. A 30 second dark-field video (18 frames per second) was immediately recorded via StreamPix 7. Multiple features of the swim behavior were then analyzed using CeleST (Restif *et al.* 2014). Graphpad Prism 6 was used for data plotting and statistics.

### Crawling assay

We washed ~30 day-1 adult hermaphrodites into M9 buffer (containing 0.2% BSA to prevent worms sticking to plastic tips) via low speed centrifuging. We transferred these worms (in 20 μl volume) to a NGM agar plate (60 mm). After the liquid was completely dried and most animals were separated from each other, we started a 30 second video recording (20 frames per second). The video was processed on ImageJ and analyzed via wrMTrck plugin (Nussbaum-Krammer *et al.* 2015). The crawling paths were generated in ImageJ and enhanced with Photoshop.

### Age pigment assay

Age pigments of day-5 adult hermaphrodite (Gerstbrein *et al.* 2005) were captured via Zeiss LSM510 Meta Confocal Laser Scanning Microscope (excitation: Water cooled Argon laser at 364nm; emission: 380nm–420nm). The auto-fluorescence intensity was quantified in ImageJ.

### Lifespan

Synchronized worms were picked at the L4 stage, and fed with OP50-1 bacteria on a 35mm NGM agar plate (12 worms per plate, ~100 animals initiating each trial). Before the end of reproductive phase, animals were transferred into a new plate every two days to keep adults separated from progeny. Immobile animals without any response to touch were counted as dead; bagged worms were also counted as deaths; animals crawling off the NGM agar were counted as lost and were excluded from analysis.

## RESULTS

### *pals-22* mutants show a transgene silencing phenotype

Based on our long-standing interest in studying the regulation of pan-neuronal gene expression (Stefanakis *et al.* 2015), we sought to use genetic mutant screens to isolate factors that control the expression of pan-neuronally expressed reporter transgenes. One screen that we undertook used a regulatory element from the pan-neuronally expressed *ric-19* locus, fused to *gfp* (*otIs381[ric-19^prom6^::NLS::gfp]*)(Stefanakis *et al.* 2015). We identified three independent mutant alleles, *ot723, ot810* and *ot811*, in which expression of *ric-19^prom6^::NLS::gfp* was reduced throughout the nervous system in all animals examined (**Fig. 1A, B**). Using our previously described whole-genome sequencing and mapping pipeline (Doitsidou *et al.* 2010; Minevich *et al.* 2012), we found that all three mutations affect the same locus, *C29F9.1* (**Fig. 2A, B**), which we named *pals-22* for reasons that we explain further below. The *ric-19^prom6^::NLS::gfp* expression defect of *pals-22(ot811)* can be rescued by a fosmid (WRM0616DC09) encompassing the *pals-22* locus plus neighboring genes as well as a genomic fragment that only contains the *pals-22* locus (791 bp upstream of the start codon to the stop codon and its 3’UTR)(**Fig. 2C**). Both *ot810* and *ot811* alleles carry early nonsense mutations and are therefore predicted to be null alleles (**Fig. 2B**).

**Fig. 1:**
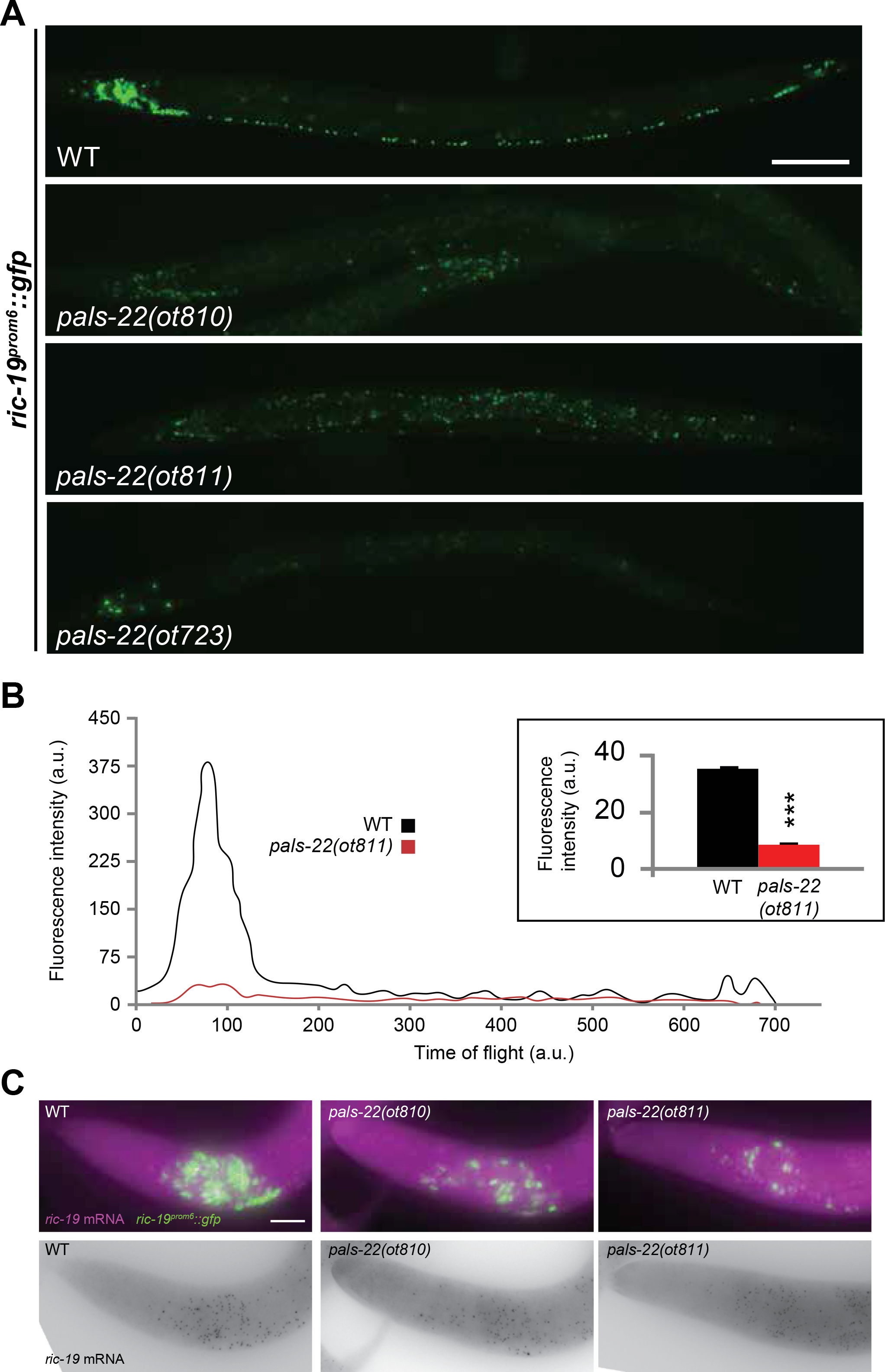
Loss of pan-neuronal reporter gene expression in *pals-22* mutants. **(A)** The *ric-19^prom6^::NLS::gfp* transcriptional reporter is brightly expressed in all neurons in wild-type N2 worms, with a nuclear localization. In *pals-22(ot723), pals-22(ot810)* and *pals-22(ot811)* mutants, expression is reduced throughout all the nervous system. All images correspond to L4 worms. Scale bar represents 50 μm **(B)** Fluorescence profiles of a representative individual day 1 adult worm expressing *ric-19^prom6^::NLS::gfp* in wild-type (black line) or *pals-22(ot811)* mutant background (red line) obtained with a COPAS FP-250 system. Time of flight indicates worm length, with lower values corresponding to the head of the worms. a.u., arbitrary units. Inset bar graph displays quantification of total fluorescence intensity averaged over 500 animals analyzed by COPAS. The data are presented as mean + SEM. Unpaired *t*-tests were performed for *pals-22(ot811)* compared to WT; \*\*\**p* < 0.001. **(C)** Wild-type (left), *pals-22(ot810)* (middle) and *pals-22(ot811)* (right) images of the anterior part of L3 worms showing equal *ric-19* mRNA levels in control and mutant worms as assessed by single molecule fluorescence *in situ* hybridization. Individual transcripts shown as purple dots in top and as black dots in bottom panels. GFP expression of the reporter transgene is shown in green. At least 20 animals examined for each genotype displayed indistinguishable staining. Scale bar 10 μm.

**Fig. 2:**
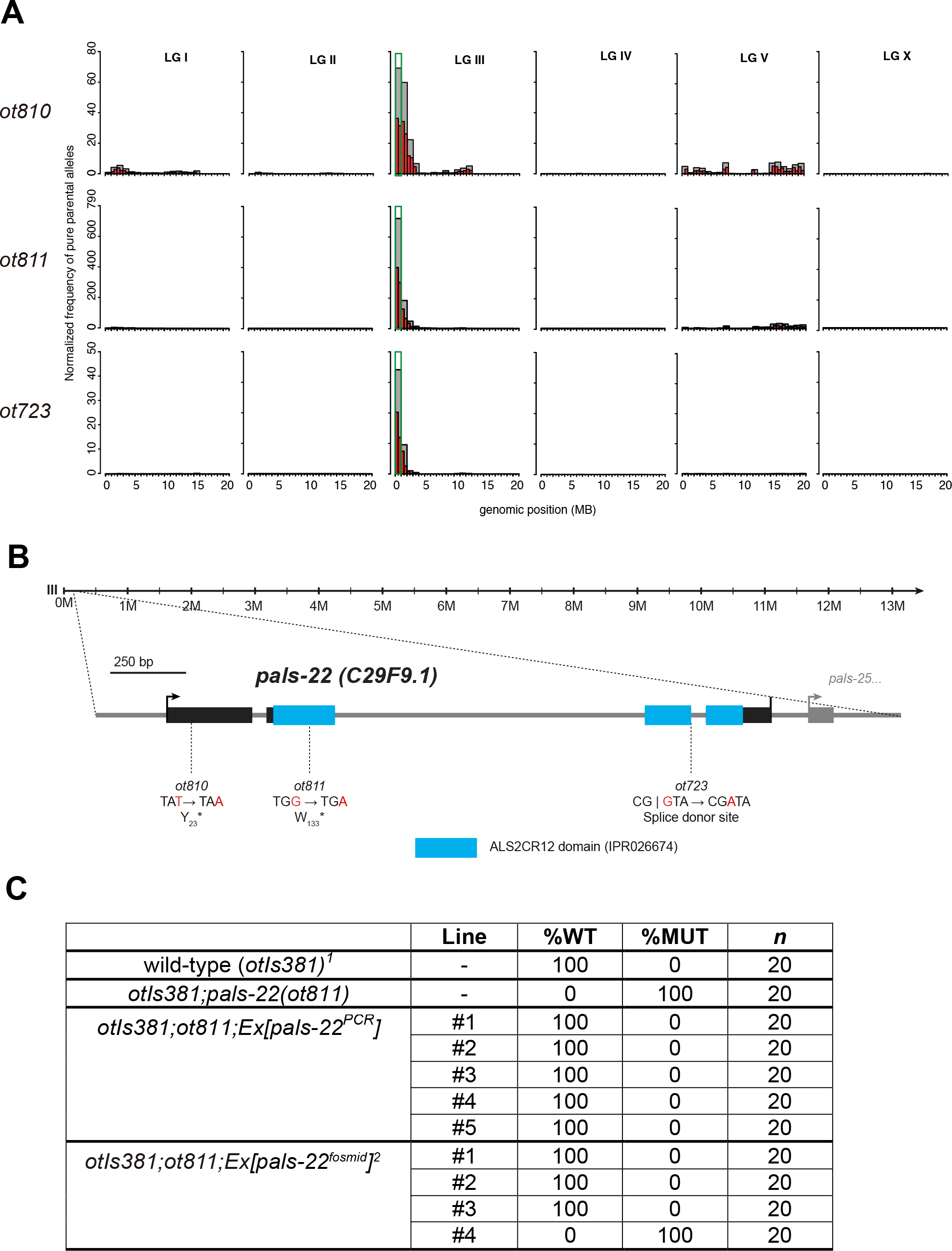
*pals-22* codes for a protein with an ALS2CR12 domain. **(A)** Hawaiian single nucleotide polymorphism (SNP) mapping plots obtained from whole-genome sequencing of the following mutant alleles: *pals-22(ot810)* (top row), *pals-22(ot811)* (middle row), and *pals-22(ot723)* (bottom row). **(B)** Schematic of *pals-22* gene locus depicting its ALS2CR12 domain and mutant allele annotation. **(C)** *pals-22* rescue data. %WT = % percent animals that express the *ric-19* reporter strongly (as in wild-type animals). %MUT = % percent animals that express the *ric-19* reporter more weakly than wild-type.^*1*^*otIs381* = *ric-19^prom6^::NLS::gfp.^2^*WRM0616DC09 fosmid.

*pals-22* mutants display reduced GFP expression of two separate *ric-19^prom6^::NLS::gfp* integrated reporter transgenes (**Table 2**). However, single molecule fluorescence *in situ* hybridization (smFISH) against endogenous *ric-19* transcripts failed to detect effects on the endogenous *ric-19* expression (**Fig. 1C**). Since the two *ric-19* reporter transgenes that are affected by *pals-22* are repetitive, “simple” arrays, we considered the possibility that *pals-22* may encode a transgene silencing activity. To test this notion, we examined the expression of a wide range of reporter transgenes in *pals-22*-deficient mutants (summarized in **Table 2**). Six additional simple arrays with widely different cellular specificities of expression (pan-neuronal, dopaminergic neurons, ventral cord motorneurons, muscle) are also silenced in *pals-22* mutants (**Table 2, Fig. 3**). Two of these arrays, *myo-3::gfp* (*ccIs4251*), and *cat-2::gfp (otIs251)* were previously shown to be silenced by loss of *tam-1*, a “classic” transgene silencer mutation (Hsieh *et al.* 1999)(M. D. and O. H. unpublished data)(**Fig. 3**). We quantified the magnitude of the *pals-22(ot811)* effect on expression of simple array reporters by acquiring fluorescence intensity information from a synchronized worm population of worms using a COPAS FP-250 system (Union Biometrica; “worm sorter”). At the L4 larval stage, we observed a 76% reduction in green fluorescence intensity for *ric-19^prom6^::NLS::gfp*, 66% reduction for *myo-3::gfp*, 32% reduction for *cat-2::gfp* and 42% reduction in red fluorescence intensity for *rab-3^prom1^::NLS::rfp* (**Fig. 1B; Fig. 3B, D, E**).

**Fig. 3:**
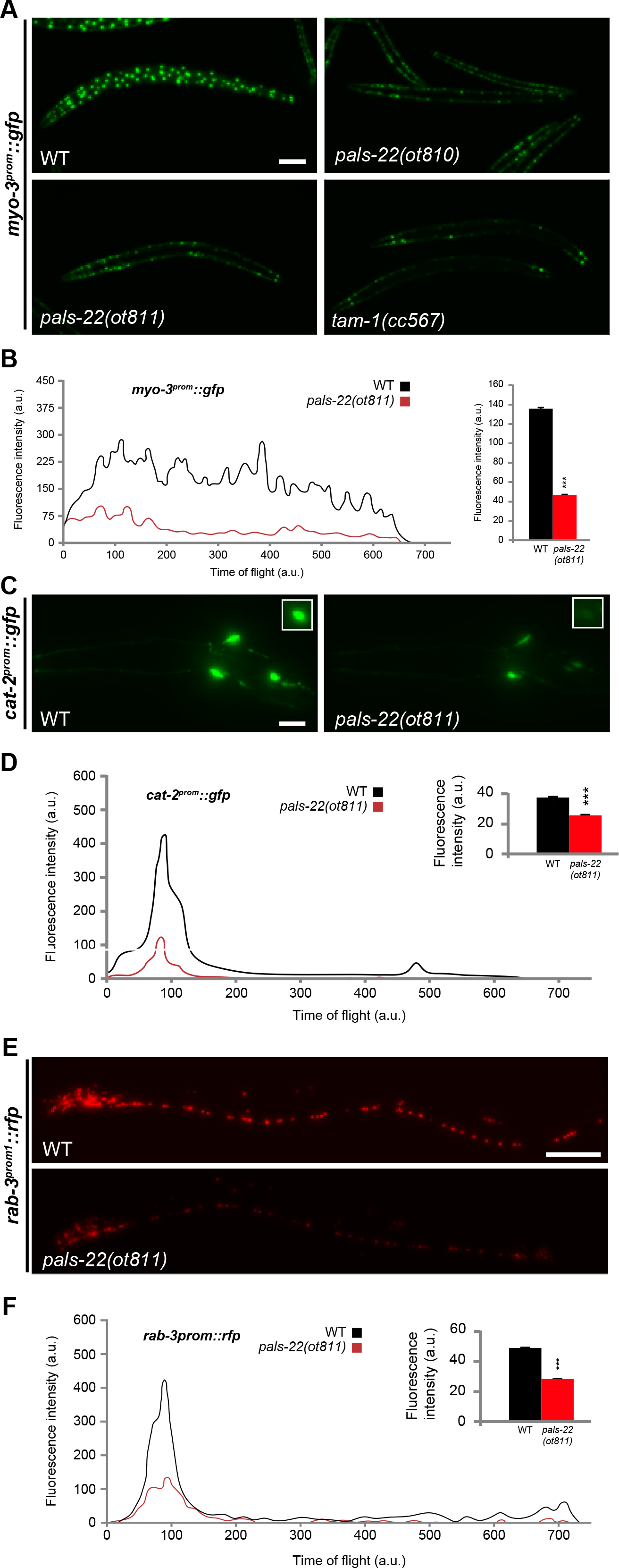
*pals-22* mutants show silencing of several multicopy arrays. **(A)** The *myo-3^prom^::gfp* transcriptional reporter is brightly expressed in all body muscles in wild-type N2 worms, with a combination of mitochondrial and nuclear localization. In *pals-22(ot810), pals-22(ot811)* and *tam-1(cc567)* mutants a generalized reduction in GFP fluorescence is observed. All images correspond to L4 worms. Scale bar represents 50 μm. **(B)** Fluorescence profile of a representative individual day 1 adult worm expressing *myo-3^prom^::gfp* in wild-type (black line) or *pals-22(ot811)* mutant background (red line) obtained with a COPAS FP-250 system. Time of flight indicates worm length, with lower values corresponding to the head of the worms. a.u., arbitrary units. Inset bar graph displays quantification of total fluorescence intensity averaged over 1000 animals analyzed by COPAS. The data are presented as mean + SEM. Unpaired *t*-tests were performed for *pals-22(ot811)* compared to WT; \*\*\**p* < 0.001. **(C)** GFP images showing silencing of *cat-2^prom^::gfp* expression in the head of *pals-22(ot811)* mutants (right) compared to wild-type L4 worms (left). The *cat-2^prom^::gfp* transcriptional reporter is expressed in all dopamine neurons, CEPD, CEPV and ADE in the head, and PDE in the posterior mid-body (top right insets). Scale bar represents 10 μm. **(D)** Fluorescence profile of a representative individual day 1 adult worms expressing *cat-2^prom^::gfp* in wild-type (black line) or *pals-22(ot811)* mutant background (red line) obtained with a COPAS FP-250 system. Time of flight indicates worm length, with lower values corresponding to the head of the worms. a.u., arbitrary units. Inset bar graph displays quantification of total fluorescence intensity averaged over 1000 animals analyzed by COPAS. The data are presented as mean + SEM. Unpaired *t*-tests were performed for *pals-22(ot811)* compared to WT; \*\*\**p* < 0.001. **(E)** RFP images showing silencing of *rab-3^prom1^::rfp* expression in the head of *pals-22(ot811)* mutants (right) compared to wild-type L4 worms (left). **(F)** Fluorescence profile of a representative individual day 1 adult worms expressing *rab-3^prom1^::rfp* in wild-type (black line) or *pals-22(ot811)* mutant background (red line) obtained with a COPAS FP-250 system. Time of flight indicates worm length, with lower values corresponding to the head of the worms. a.u., arbitrary units. Inset bar graph displays quantification of total fluorescence intensity averaged over 1000 animals analyzed by COPAS. The data are presented as mean + SEM. Unpaired *t*-tests were performed for *pals-22(ot811)* compared to WT; \*\*\**p* < 0.001.

**Table 2:** Pals-22 effects on transgene reporter expression.

| Array type | Construct | Expression pattern | Relative intensities of reporters |  |
| --- | --- | --- | --- | --- |
|  |  |  | wild-type | *pals-22*<sup>1</sup> |
| Simple array | *ric-19p6::gfp*<sup>2</sup> | Pan-neuronal | +++ | + |
|  | *myo-3p::gfp* | Body-wall muscle | ++++ | + |
|  | *cat-2p::gfp* | Dopaminergic neurons | ++++ | ++ |
|  | *rab-3p1::rfp* | Pan-neuronal | +++ | + |
|  | *unc-3p::mCherry* | Cholinergic neurons | ++++ | ++ |
|  | *tdc-1p::yfp* | RIC and RIM neurons | ++ | + |
|  | *ric-4p26::yfp*<sup>3</sup> | Neuronal | ++++ | + |
| Complex array | *unc-11p8::gfp* | Pan-neuronal | +++ | + |
|  | *cho-1fosmid::yfp* | Cholinergic neurons | ++ | ++ |
|  | *ric-4fosmid::yfp* | Pan-neuronal | ++ | ++ |
| Single copy | *lin-4p::yfp* | Ubiquitous | ++ | ++ |
|  | *myo-3p::mRuby* | Pharyngeal muscle | ++ | ++ |
| None<br>(Endogenous tag) | *che-1::gfp* | ASE neurons | + | + |

<sup>1</sup> *pals-22(ot811)* mutant background.
<sup>2</sup> *otIs381* and *otIs380* strains (independently integrated lines).
<sup>3</sup> Extrachromosomal array.

We also analyzed the expression of complex array (tandemly repeated transgenes with a less repetitive structure) and single copy reporter transgenes in *pals-22* mutants (summarized in **Table 2**). One complex array transgene was silenced (*unc-11p8::gfp*, **Table 2**) but others were not (*ric-4^fosmid^::yfp* and *cho-1^fosmid^::yfp*, **Table 2, Fig. S1A**). Perhaps the much smaller reporter fragment *unc-11p8::gfp* generated more repetitive structures even in the context of “complex” arrays than the larger fosmid-based reporters. Importantly, single copy insertions or endogenously tagged genes are not affected by mutations in *pals-22* (**Fig. S1B, C**).

Similar to previously characterized transgene silencing mutants, such as *tam-1* (Hsieh *et al.* 1999), the reduction of reporter expression is temperature sensitive. However, the direction of the sensitivity is inverted as compared to the *tam-1* case: the decrease in *ric-19^prom6^::NLS::gfp* expression is most pronounced at 15°C, while the effect is milder at 25°C (data not shown). We also found stage dependent variability: the decrease in transgene reporter expression is stronger as the animals develop, from mild differences in expression in early stages of development to more obvious defects at later stages (**Fig. S2**).

### PALS-22 is a broadly expressed, cytoplasmic protein

To analyze the expression pattern of *pals-22* we fused the entire locus to *gfp* (including 791 bp of 5’ sequences and all exons and introns; **Fig. 4A**). This reporter construct fully rescues the transgene silencing phenotype of *pals-22(ot811)* mutants (**Fig. 4D**). Expression is observed widely across different tissues: nervous system (pan-neuronal expression), body wall and pharyngeal muscle, gut, seam cells, as well as the male tail (**Fig. 4B, C**). A similar expression pattern is observed with a transcriptional reporter (791 bp of its upstream region fused to GFP; **Fig. 4C**). The rescuing, translational reporter transgene revealed a strong, if not exclusive enrichment in the cytoplasm and appears to be excluded from the nucleus in most tissues (**Fig. 4B**). However, we cannot exclude the possibility that the nucleus contains low amount of functional PALS-22 protein.

**Fig. 4:**
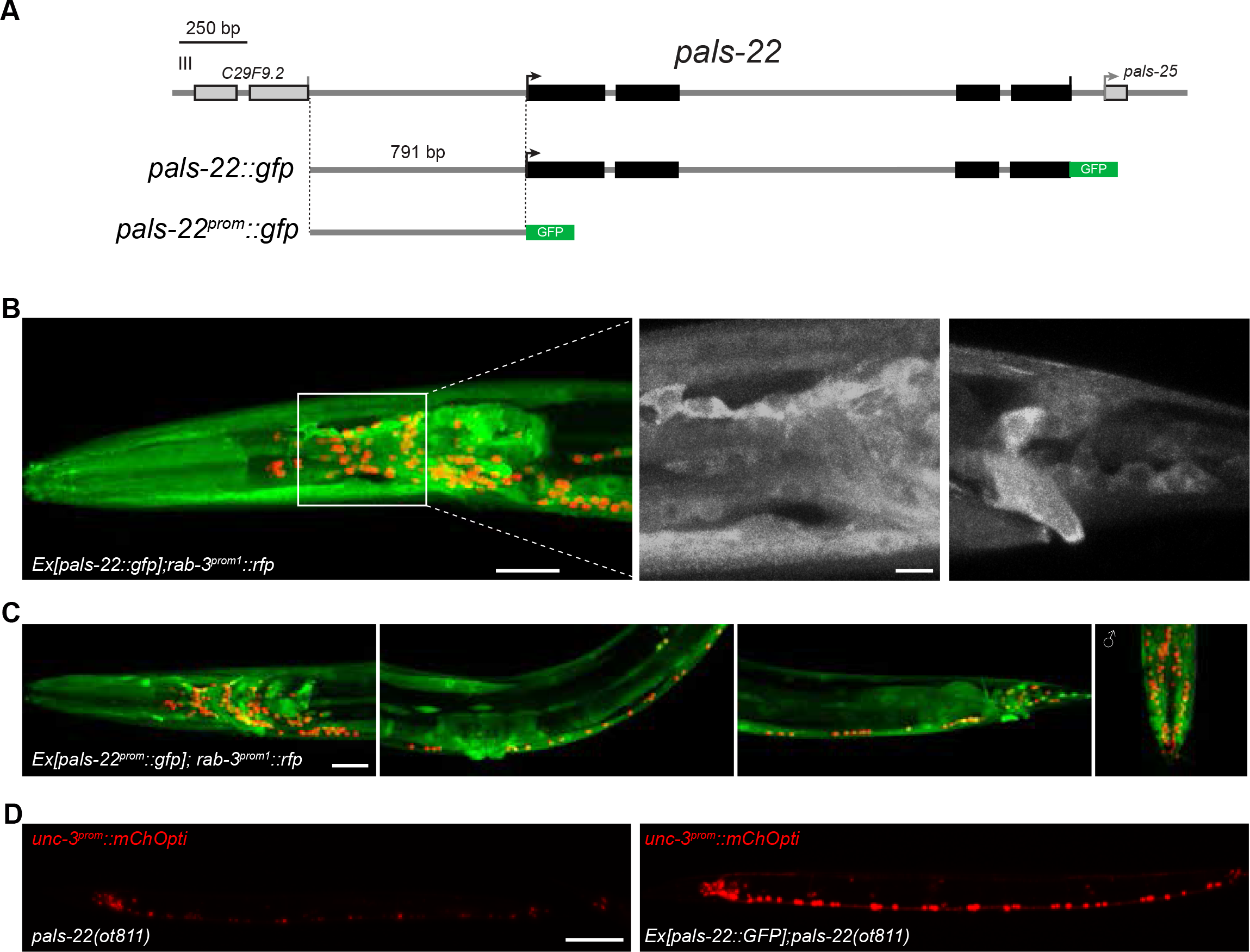
PALS-22 is a broadly expressed, cytoplasmic protein. **(A)** Schematic of *pals-22* transcriptional and translational GFP reporters. The *gfp* reporter is followed by the 3’UTR from the *unc-54* gene. **(B)** Expression of the *pals-22::gfp* translational reporter (*otEx7037*) is shown in green in the head (left) in a representative hermaphrodite L4 worm. High magnification images in black and white for the head (middle) and tail (right) show PALS-22 cytoplasmic localization. Nuclear pan-neuronal *rab-3^prom1^::rfp* expression is shown in red on the background. Two distinct extrachromosomal arrays show the same pattern. **(C)** Expression of the *pals-22^prom^::gfp* transcriptional reporter (*otEx7036*) is shown in green in the head (left panel), mid-body (second panel), and tail (third panel) in a representative hermaphrodite L4 worm; and in the male tail (right panel). Nuclear pan-neuronal *rab-3^prom1^::rfp* expression is shown in red on the background. Three distinct extrachromosomal arrays show the same pattern. **(D)** PALS-22::GFP (right) rescues the expression of *unc-3^prom^::mChOpti* in *pals-22(ot811)* mutants (left). At least 50 animals examined for each genotype. Scale bars represent 20 μm (B and C), 5 μm (B, high magnification images) and 50 μm (D).

### *pals-22* is a member of an unusual *C. elegans* gene family

The molecular analysis of the PALS-22 protein sequence revealed its membership in an unusual gene family. The only recognizable feature of PALS-22 is a domain termed ALS2CR12 by the InterPro (https://www.ebi.ac.uk/interpro/) (Finn *et al.* 2017) and Panther (http://www.pantherdb.org/)(Mi *et al.* 2013; Mi *et al.* 2017) databases (hence the name “PALS” for “<u>p</u>rotein containing <u>ALS</u>2CR12 domain”). This domain was first found in a human gene that constituted a potential disease locus for amyotrophic lateral sclerosis 2 (Hadano *et al.* 2001). No biochemical or cellular function has yet been assigned to the human ALS2C12 protein or any of its homologs in other organisms.

Both InterPro and Panther databases group PALS-22 within a family of ALS2CR12 domain containing proteins from different species ranging from nematodes to vertebrates (**Fig. 5A**). While there is only one ALS2CR12 domain containing protein in vertebrates such as mouse or human, the number of proteins containing this domain is strikingly expanded to a total of 39 distinct proteins in *C. elegans* (**Fig. 5A; Table 3**). *Drosophila* seems to be completely devoid of ALS2CR12 domain-containing proteins.

**Fig. 5:**
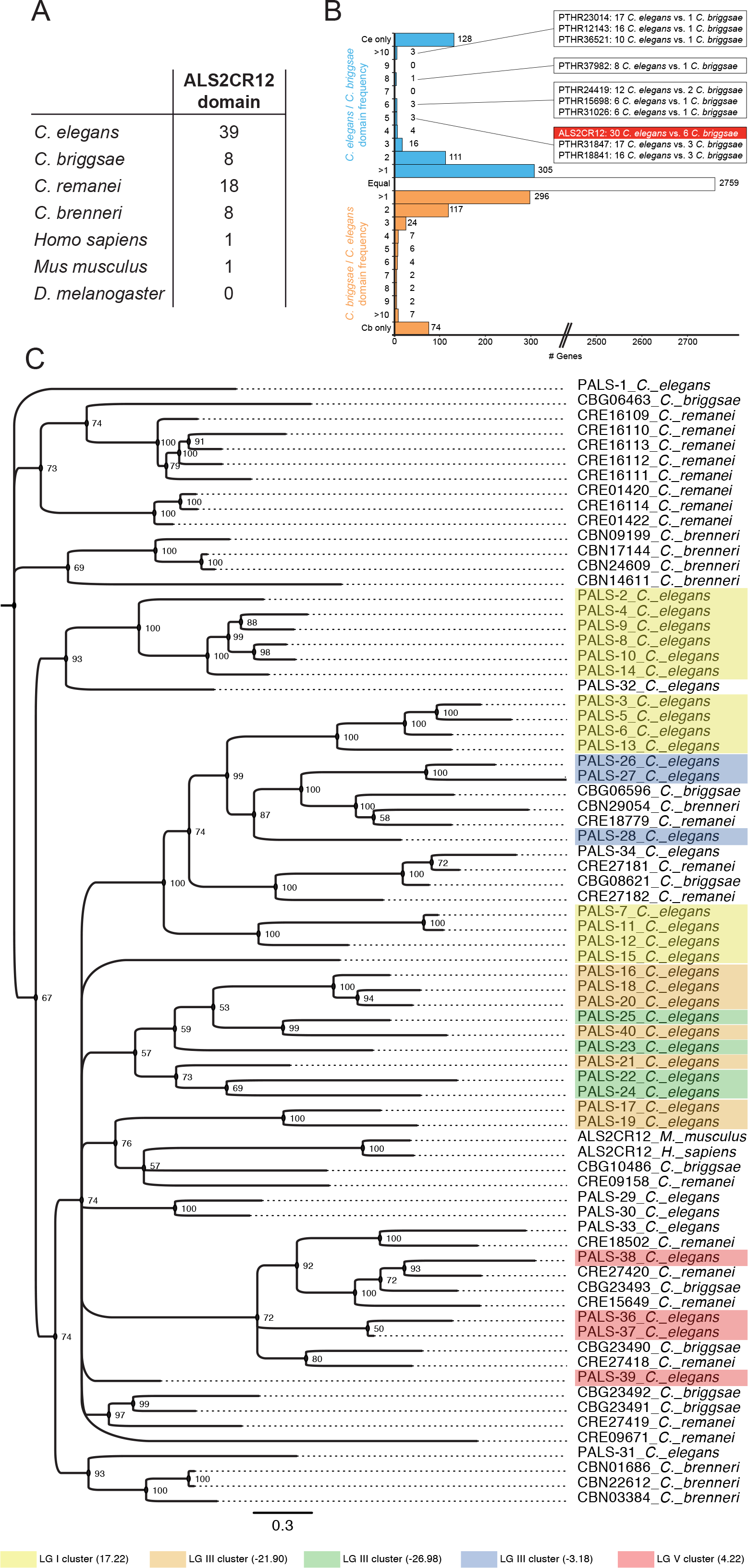
Sequence and genomic analysis of *pals* gene family in *C. elegans* and other species. **(A)** Number of genes containing the ALS2CR12 domain in their protein product as predicted by InterPro (IPR026674) and/or Panther (PTHR21707). **(B)** Graph representing the *C. elegans* to *C. briggsae* domain frequency for domains enriched in *C. elegans* (blue), *C. briggsae* to *C. elegans* ratio for domains enriched in *C. briggsae* (orange), or white for Panther protein domains predicted in the same number of genes in both species. Boxes indicate the gene counts for highly enriched domains in *C. elegans.* The number of Panther domain hits for the *C. elegans* and *C. briggsae* genome were obtained from WormBase. **(C)** Phylogram of ALS2CR12 domain containing genes, including *C. elegans* paralogs and orthologs from *C. briggsae, C. remanei, C. brenneri*, human and mouse. Node values indicate posterior probabilities for each split as percent. The scale bar indicates average branch length measured in expected substitutions per site.

**Table 3:**
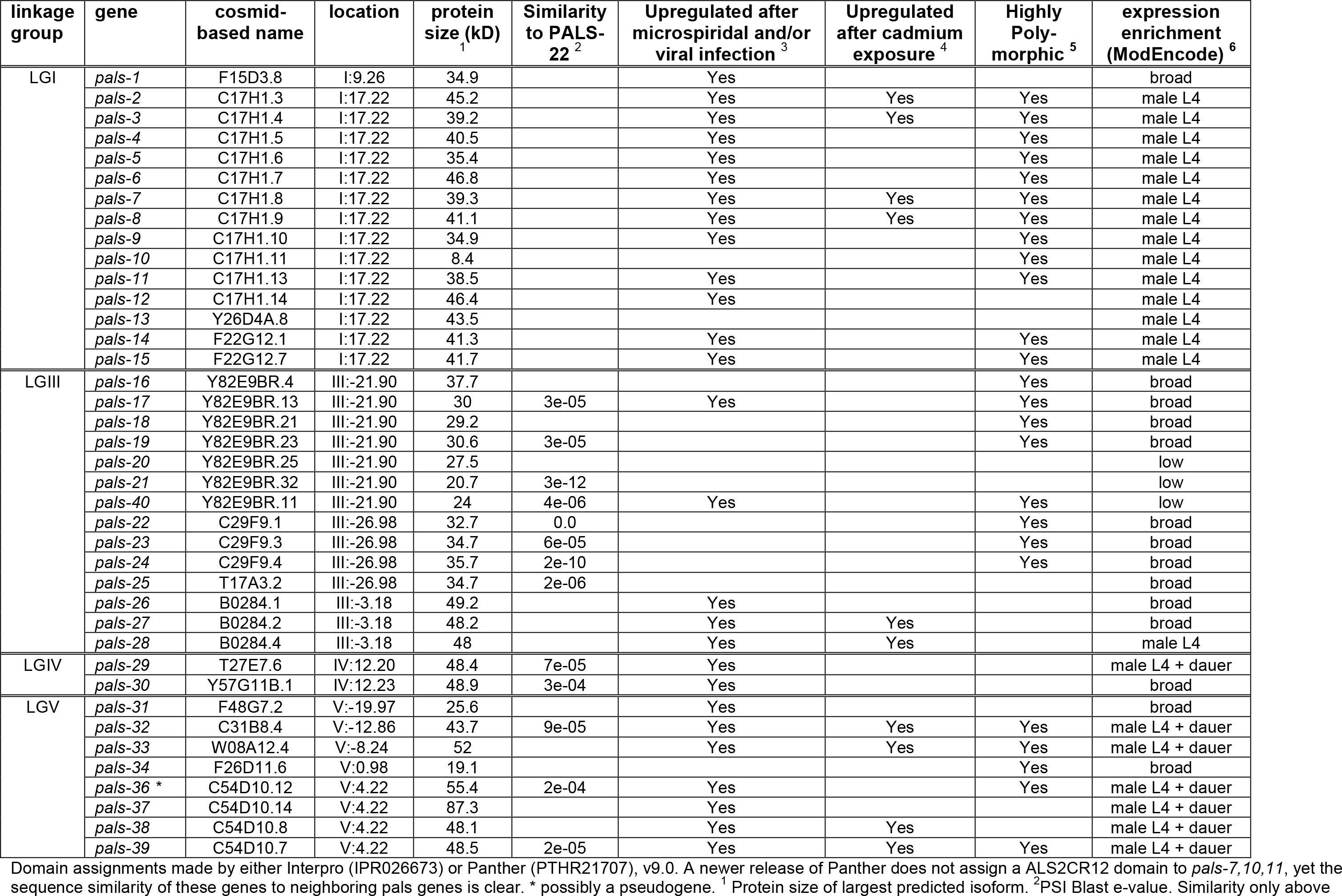
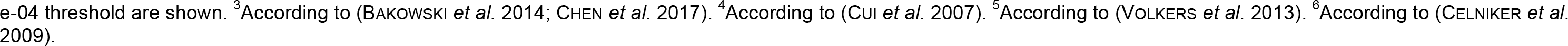
List of all pals genes

| linkage group | gene | cosmid-based name | location | protein size (kD)<br><sup>1</sup> | Similarity to PALS-22<sup>2</sup> | Upregulated after microspiridal and/or viral infection<sup>3</sup> | Upregulated after cadmium exposure<sup>4</sup> | Highly Poly-morphic<sup>5</sup> | expression enrichment (ModEncode)<sup>6</sup> |
| --- | --- | --- | --- | --- | --- | --- | --- | --- | --- |
| LGI | *pals-1* | F15D3.8 | I:9.26 | 34.9 |  | Yes |  |  | broad |
|  | *pals-2* | C17H1.3 | I:17.22 | 45.2 |  | Yes | Yes | Yes | male L4 |
|  | *pals-3* | C17H1.4 | I:17.22 | 39.2 |  | Yes | Yes | Yes | male L4 |
|  | *pals-4* | C17H1.5 | I:17.22 | 40.5 |  | Yes |  | Yes | male L4 |
|  | *pals-5* | C17H1.6 | I:17.22 | 35.4 |  | Yes |  | Yes | male L4 |
|  | *pals-6* | C17H1.7 | I:17.22 | 46.8 |  | Yes |  | Yes | male L4 |
|  | *pals-7* | C17H1.8 | I:17.22 | 39.3 |  | Yes | Yes | Yes | male L4 |
|  | *pals-8* | C17H1.9 | I:17.22 | 41.1 |  | Yes | Yes | Yes | male L4 |
|  | *pals-9* | C17H1.10 | I:17.22 | 34.9 |  | Yes |  | Yes | male L4 |
|  | *pals-10* | C17H1.11 | I:17.22 | 8.4 |  |  |  | Yes | male L4 |
|  | *pals-11* | C17H1.13 | I:17.22 | 38.5 |  | Yes |  | Yes | male L4 |
|  | *pals-12* | C17H1.14 | I:17.22 | 46.4 |  | Yes |  |  | male L4 |
|  | *pals-13* | Y26D4A.8 | I:17.22 | 43.5 |  |  |  |  | male L4 |
|  | *pals-14* | F22G12.1 | I:17.22 | 41.3 |  | Yes |  | Yes | male L4 |
|  | *pals-15* | F22G12.7 | I:17.22 | 41.7 |  | Yes |  | Yes | male L4 |
| LGIII | *pals-16* | Y82E9BR.4 | III:-21.90 | 37.7 |  |  |  | Yes | broad |
|  | *pals-17* | Y82E9BR.13 | III:-21.90 | 30 | 3e-05 | Yes |  | Yes | broad |
|  | *pals-18* | Y82E9BR.21 | III:-21.90 | 29.2 |  |  |  | Yes | broad |
|  | *pals-19* | Y82E9BR.23 | III:-21.90 | 30.6 | 3e-05 |  |  | Yes | broad |
|  | *pals-20* | Y82E9BR.25 | III:-21.90 | 27.5 |  |  |  |  | low |
|  | *pals-21* | Y82E9BR.32 | III:-21.90 | 20.7 | 3e-12 |  |  |  | low |
|  | *pals-40* | Y82E9BR.11 | III:-21.90 | 24 | 4e-06 | Yes |  | Yes | low |
|  | *pals-22* | C29F9.1 | III:-26.98 | 32.7 | 0.0 |  |  | Yes | broad |
|  | *pals-23* | C29F9.3 | III:-26.98 | 34.7 | 6e-05 |  |  | Yes | broad |
|  | *pals-24* | C29F9.4 | III:-26.98 | 35.7 | 2e-10 |  |  | Yes | broad |
|  | *pals-25* | T17A3.2 | III:-26.98 | 34.7 | 2e-06 |  |  |  | broad |
|  | *pals-26* | B0284.1 | III:-3.18 | 49.2 |  | Yes |  |  | broad |
|  | *pals-27* | B0284.2 | III:-3.18 | 48.2 |  | Yes | Yes |  | broad |
|  | *pals-28* | B0284.4 | III:-3.18 | 48 |  | Yes | Yes |  | male L4 |
| LGIV | *pals-29* | T27E7.6 | IV:12.20 | 48.4 | 7e-05 | Yes |  |  | male L4 + dauer |
|  | *pals-30* | Y57G11B.1 | IV:12.23 | 48.9 | 3e-04 | Yes |  |  | broad |
| LGV | *pals-31* | F48G7.2 | V:-19.97 | 25.6 |  | Yes |  |  | broad |
|  | *pals-32* | C31B8.4 | V:-12.86 | 43.7 | 9e-05 | Yes | Yes | Yes | male L4 + dauer |
|  | *pals-33* | W08A12.4 | V:-8.24 | 52 |  | Yes | Yes | Yes | male L4 + dauer |
|  | *pals-34* | F26D11.6 | V:0.98 | 19.1 |  |  |  | Yes | broad |
|  | *pals-36 \** | C54D10.12 | V:4.22 | 55.4 | 2e-04 | Yes |  | Yes | male L4 + dauer |
|  | *pals-37* | C54D10.14 | V:4.22 | 87.3 |  | Yes |  |  | male L4 + dauer |
|  | *pals-38* | C54D10.8 | V:4.22 | 48.1 |  | Yes | Yes |  | male L4 + dauer |
|  | *pals-39* | C54D10.7 | V:4.22 | 48.5 | 2e-05 | Yes | Yes | Yes | male L4 + dauer |

Domain assignments made by either Interpro (IPR026673) or Panther (PTHR21707), v9.0. A newer release of Panther does not assign a ALS2CR12 domain to *pals-7,10,11*, yet the sequence similarity of these genes to neighboring *pals* genes is clear. \* possibly a pseudogene. <sup>1</sup> Protein size of largest predicted isoform. <sup>2</sup>PSI Blast e-value. Similarity only above
e-04 threshold are shown. <sup>3</sup>According to (BAKOWSKI *et al.* 2014; CHEN *et al.* 2017). <sup>4</sup>According to (CUI *et al.* 2007). <sup>5</sup>According to (VOLKERS *et al.* 2013). <sup>6</sup>According to (CELNIKER *et al.* 2009).

The 39 *C. elegans* PALS proteins are very divergent from one another, as reflected in **Table 3** with similarity scores of PALS-22 compared to other PALS proteins. Besides poor paralogy, there is also poor orthology. For example, BLAST searches with PALS-22 picks up no sequence ortholog in *C. briggsae*, and the best homolog from another species (*C. brenneri, CBN22612*) is less similar to PALS-22 than some of the *C. elegans* PALS-22 paralogs. The divergence of *C. elegans pals* genes is also illustrated by the genome sequence of wild isolates of *C. elegans* in which many *pals* genes display an above-average accumulation of polymorphisms (Volkers *et al.* 2013)(**Table 3**).

The expansion of *C. elegans* ALS2CR12 domain-containing proteins appears to be nematode species-specific, as *C. briggsae* only contains 8 predicted proteins with an ALS2CR12 domain; other nematodes also contain significantly less ALS2CR12 domain proteins (**Fig. 5A**). Such *C. elegans-specific* gene expansion is highly unusual, as shown in **Fig. 5B**. Among 3874 Panther protein domains analyzed, 2759 (71.2%) are present in the same number of genes in both *C. elegans* and *C. briggsae*, 128 (3.3%) domains are present only in *C. elegans*, while 74 (1.9%) are present only in *C. briggsae.* Most of the remaining domains are only slightly enriched in one species or the other. Just 10 domains (0.3%) are enriched five times or more in *C. elegans* versus *C. briggsae* (**Fig. 5B**). Most of these domain families contain uncharacterized genes, even though many of them contain human and vertebrate orthologs. Among them, the ALS2CR12 domain has not only a noteworthy enrichment, but it also constitutes the family with the largest absolute number of genes (**Fig. 5B**).

As perhaps expected from a species-specific expansion, the ALS2CR12 domain-encoding *pals* genes are genomically clustered (**Fig. 6**). 14 genes are clustered in chromosome I (position I: 17.22 cM); while three clusters with 7 genes (III: -21.90 cM), 4 genes (III: -26.98 cM) and 3 genes (III: -3.18 cM) are present in chromosome III; lastly 4 genes are clustered in chromosome V (V: 4.22) (**Fig. 6; Table 3**). Taking into account the *C.* elegans-specific family expansion it is not surprising to find a total lack of conservation in the regions encompassing most of the *pals* clusters (**Fig. 6**). Only one cluster on chromosome V shows some degree of conservation among other nematode species. Genes surrounding these regions are conserved, suggesting recent gene duplications in the non-conserved areas. The low conservation region in cluster III: - 21.90 cM, contains many additional, non-conserved genes that are largely expanded in a nematode-specific manner, namely the previously analyzed *fbxa* genes (Thomas 2006).

**Fig. 6:**
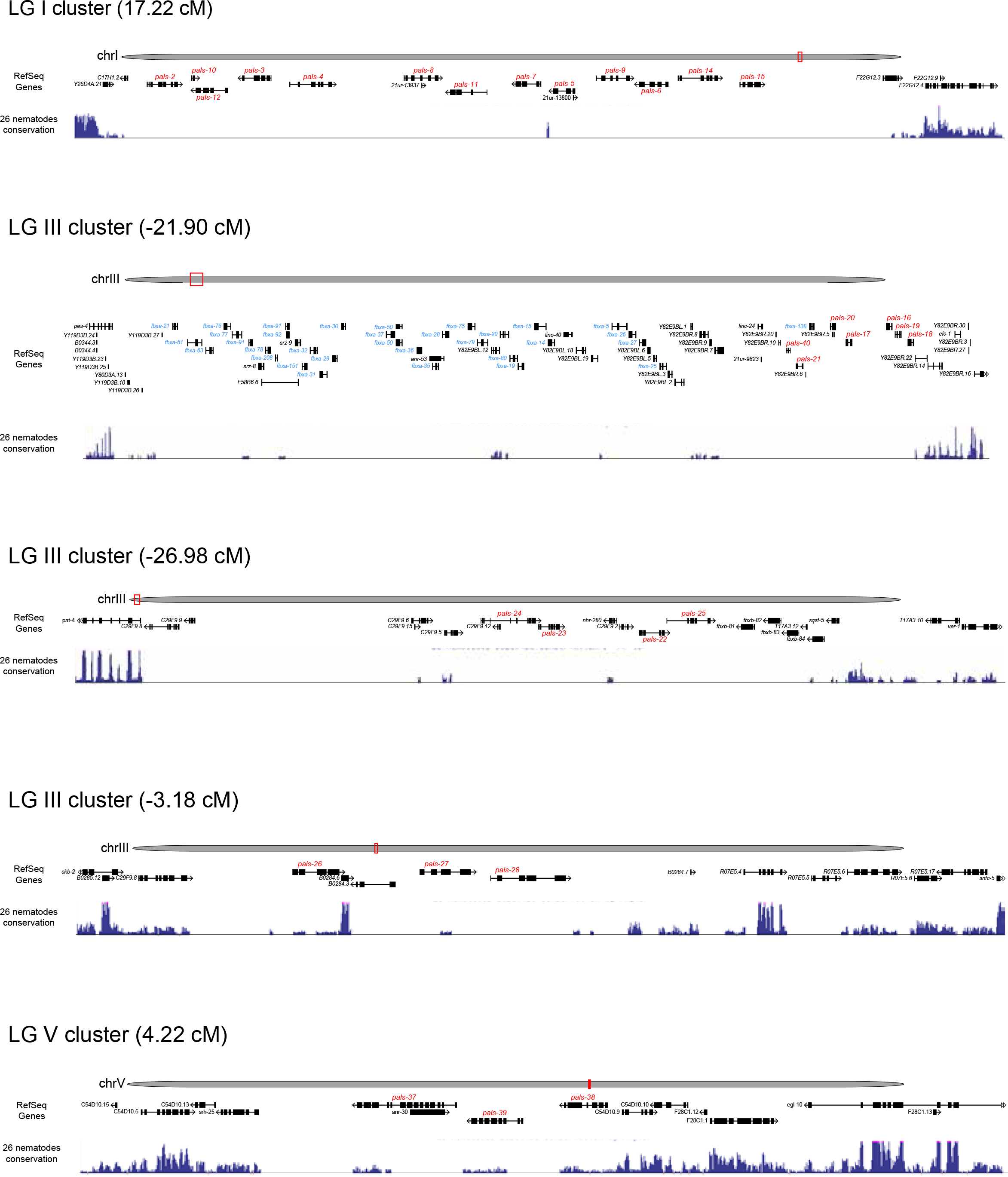
*C. elegans* pals genes are clustered and these clusters are poorly conserved. Schematics of different *C. elegans* genomic regions adapted from the UCSC Genome Browser (https://genome.ucsc.edu/). For each panel is represented, from top to bottom:*C elegans* genome assembly (RefSeq genes) and conservation of 26 distinct nematode species (Basewise Conservation by PhyloP track). One isoform per gene is shown. *pals* genes are indicated in red, *fbxa* genes in blue. The following regions are shown: Chromosome I, cluster at position 17.22 cM, chrl: 13,099,564-13,160,497 bp. Chromosome Ill, cluster at position -21.90 cM, chrIII: 1,215,665-1,423,550 bp. Chromosome III, cluster at position -26.98 cM, chrIII: 89,907-159,962 bp. Chromosome III, cluster at position -3.18 cM, chrIII: 4,368,479-4,405,090 bp. Chromosome V, cluster at position 4.22 cM, chrV: 12,427,096-12,462,015 bp.

Local gene duplications seem a plausible mechanism for the origin of the expanded *C. elegans pals* gene family. Consequently, we reasoned that *pals* genes within the same cluster should be more similar among each other than to other *pals.* In order to explore this possibility, we built a phylogenetic tree to visualize the phylogenetic relationships between *pals* genes (**Fig. 5C**). We included all *C. elegans pals* genes plus orthologs from *C. briggsae, C. remanei, C. brenneri*, mouse and human (based on presence of InterPro IPR026674). As expected, *pals* genes within the same cluster have a closer phylogenetic relationship, suggesting a shared origin.

According to modENCODE expression data (Celniker *et al.* 2009), most of the genes within each cluster are expressed in the same stage (*e.g.* all genes clustered in chromosome I are only found in L4 males), suggesting related functions (**Table 3**). Perhaps most intriguingly, though, the majority of *pals* genes become upregulated upon exposure to specific pathogens, specifically the exposure to intracellular fungal pathogen (microsporidia) or by viral infection (Bakowski *et al.* 2014; Chen *et al.* 2017). Induction of *pals* gene expression is also observed upon various other environmental insults (exposure to toxic compounds)(Cui *et al.* 2007)(summarized in **Table 3**). Several *fbxa* genes present in the low conservation region in cluster III: -21.90 cM, also become upregulated upon exposure to microsporidia or by viral infection (Bakowski *et al.* 2014; Chen *et al.* 2017).

### Somatic transgene silencing in *pals-22* mutants requires *rde-4*-dependent small RNAs

We examined whether two *pals-22* paralogs, *pals-19* and *pals-25* may also be involved in transgene silencing. Both genes show significant sequence similarity to *pals-22* (**Table 3**) and one (*pals-25*) is directly adjacent to *pals-22* (**Fig.6**). We tested whether two nonsense alleles generated by the Million Mutant project, *pals-19(gk16606)* and *pals-25(gk891046)* (Thompson *et al.* 2013) silence the *ric-19^prom6^::NLS::gfp* and *myo-3::gfp* multi-copy transgenes. Neither array shows obvious changes in expression in *pals-19* or *pals-25* mutant backgrounds (data not shown).

To further pursue the role of *pals-22* in transgene silencing we considered previous reports on transgene silencer mutations. Several genes with transgene silencing effects are known to be involved in modifying chromatin (Cui AND Han 2007). However, the cytoplasmic localization of PALS-22 described above argues against a direct role in controlling chromatin architecture (although, a function in the nucleus can not be ruled out). Nevertheless, we do find that *pals-22* affects transgene silencing on the transcriptional level. smFISH against *gfp* mRNA shows that silenced *gfp* transgenic arrays display a significantly reduced number of transcripts in *pals-22* mutants (**Fig. 7A**). We therefore considered the possibility that the transcriptional effects of a cytoplasmic protein on transcription may be controlled by intermediary factors. Small interfering RNAs are known to affect gene expression on the transcriptional level (Zamore *et al.* 2000; Elbashir *et al.* 2001) and through a genetic epistasis test, we asked whether *pals-22* requires small RNAs for its function. To this end, we turned to *rde-4* mutant animals. The dsRNA-binding protein RDE-4 initiates gene silencing by recruiting an endonuclease to process long dsRNA into short dsRNA and is involved in exogenous as well as endogenous RNAi pathways (Tabara *et al.* 2002; Parker *et al.* 2006; Gent *et al.* 2010; Vasale *et al.* 2010). We find that loss of *rde-4* completely suppresses the *pals-22* mutant phenotype (**Fig. 7B, C**). This result demonstrates that the gene silencing mediated by *pals-22* deficiency requires the production of small dsRNAs.

**Fig. 7:**
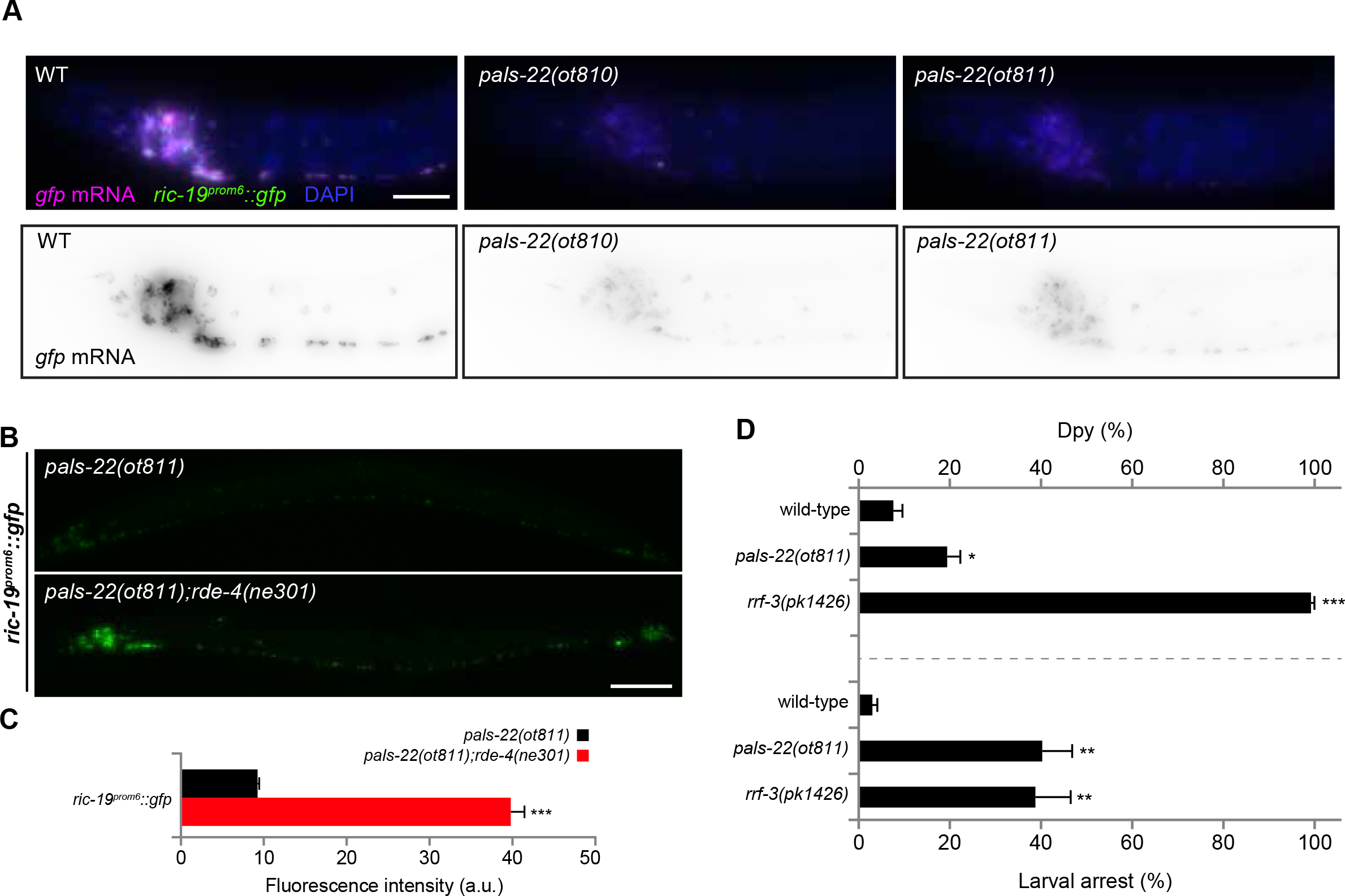
Somatic transgene silencing in *pals-22* depends on the RNAi pathway. **(A)** Wild-type (left), *pals-22(ot810)* (middle) and *pals-22(ot811)* (right) images of the anterior part of L3 worms showing reduced *gfp* mRNA levels in mutants compared to control wild-type worms as assessed by single molecule fluorescence *in situ* hybridization. lndividual transcripts shown as purple dots in top and as black dots in bottom panels. GFP expression is shown in green, DAPl staining is shown in blue. At least 20 animals examined for each genotype. **(B)** Silencing of *ric-19^prom6^::NLS::gfp* in *pals-22(ot811)* (top) is suppressed in *pals-22(ot811);rde-4(ne301)* double mutants (bottom). **(C)** Quantification of the data represented in (B). The data are presented as mean + SEM. Unpaired *t*-test, \*\*\**p* < 0.001; *n* ≥ 300-1000 for all genotypes. **(D)** Animals of the indicated genotype were grown on bacteria expressing *dpy-13* or *cel-1* dsRNA (feeding RNAi). For *dpy-13* (top), progeny were scored for the percentage of animals having a dumpy body shape (Dpy). For *cel-1* (bottom), the percentage of their progeny arresting at the L2 larval stage was determined. Wild-type *otIs381[ric-19^prom6^::NLS::gfp]* and *rrf-3(pk1426)* mutants were used as negative and positive controls. The data are presented as mean + SEM among the data collected from at least four independent experiments. Unpaired *t*-tests were performed for *pals-22(ot811)* and *rrf-3(pk1426)* compared to WT; \*\*\**p* < 0.001, \*\**p* < 0.01, \**p* < 0.05. Scale bars represent 10 μm (A) and 50 μm (B).

Transgene silencing phenotypes have been observed in mutants that affect multiple distinct small RNA pathways, and exogenous RNAi responses are often enhanced in these mutants (Simmer *et al.* 2002; Lehner *et al.* 2006; Fischer *et al.* 2013). Thus, we tested whether *pals-22(ot811)* shows an enhanced exogenous RNAi response. Using *rrf-3(pk1426)* as a positive control (Simmer *et al.* 2002), we detect enhanced *dpy-13* or *cel-1* RNAi phenotypes in *pals-22(ot811)* (dsRNA delivered by feeding; **Fig. 7D**). We conclude that *pals-22* physiological function might be related to the regulation of RNAi-dependent silencing in the cytoplasm, via a mechanism critical to its action as an anti-silencing factor.

### Locomotory and aging defects of *pals-22* mutant animals

The loss of *pals-22* has striking physiological consequences. Since casual observation of *pals-22* mutants indicates defects in locomotion, we quantified these defects, by measured swimming behavior (Restif *et al.* 2014) and crawling activity, comparing wild-type, *pals-22(ot810), pals-22(ot811)* and *pals-22(ot811)* carrying a wild-type copy of *pals-22* on an extrachromosomal array (**Fig. 8**). Day 1 adult *pals-22* mutants show a poor performance in swimming assays as evaluated by multiple parameters, including low wave initiation rate (akin to thrash speed), travel speed (distance moved over time), brush stroke area (area covered in unit time) and activity index (**Fig. 8A**). On agar plates animals also were clearly impaired, displaying significantly decreased traveling speed (**Fig. 8B**). All locomotory defects were rescued by the *pals-22(+)* extrachromosomal array.

**Fig. 8:**
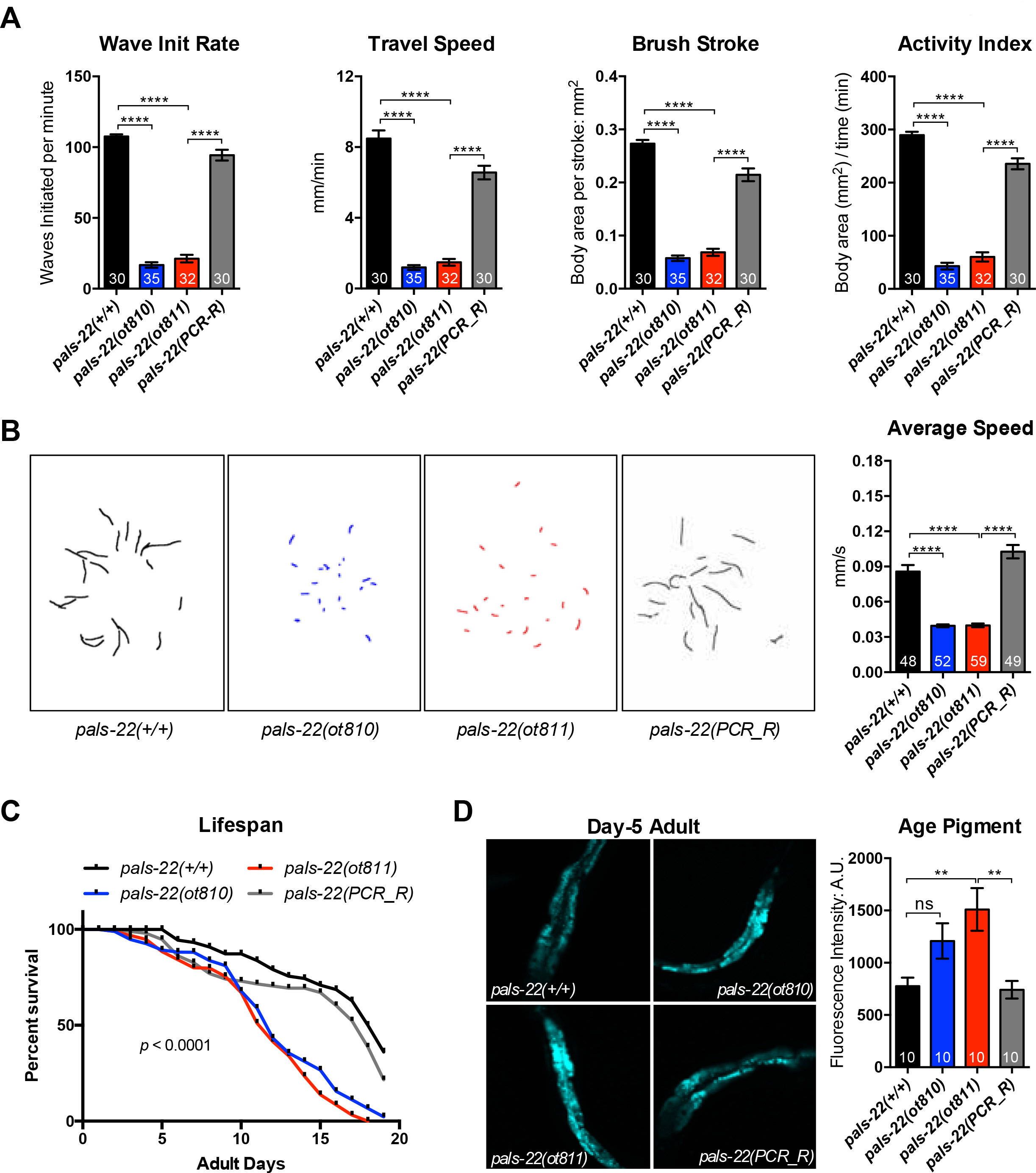
*pals-22* mutants show defective locomotion and early onset of aging traits. **(A)** Day-1 adult *pals-22* mutants exhibit defective swimming features, including decreased wave initiation rate, travel speed, brush stroke and activity index. *pals-22(PCR_R)* is the *pals-22(ot811)* mutant carrying a wild-type copy of *pals-22* on an extrachromosomal array amplified by PCR. Data shown are mean ± SEM of each parameters, n = 30-35 (number indicated in each bar) from two independent experiments, *** *p* < 0.0001 (one-way ANOVA) compared to related control. **(B)** *pals-22* mutants crawl slowly on agar plates. The left panel shows the crawling path of each animal in 30 seconds. The right panel shows the mean ± SEM of average crawling speed (millimeters per second, mm/s) for day-1 adults from two independent experiments, n = ~55 worms for each genotype, *** *p* < 0.0001 (one-way ANOVA) compared to related control. **(C)** *pals-22* mutants have shorter lifespans compared to wild type or *pals-22(PCR_R).* Survival study was initiated with 96 worms (for each genotype) at L4 stage (day-0). Data shown is one represented trial of lifespan; two additional independent trials also show similar changes of lifespan in *pals-22* mutants, *p* < 0.0001 (Log-rank test), comparing the wild type to mutants, or comparing *pals-22(PCR_R)* against *pals-22(ot811).* **(D)** *pals-22* mutants exhibit early accumulation of age pigment. The left panel shows representative pictures of age pigment (excitation: 364nm; emission: 380-420nm). The right panel shows the mean ± SEM of age pigment auto-fluorescence intensity of day-5 adults, n = 10 worms from each genotype, ns (not significant) or ** *p* < 0.01 (one-way ANOVA) compared to related control

Apart from the locomotory defects, we also noted abnormal survival of *pals-22* mutants. Using standard lifespan assays, we observed premature death of *pals-22* mutants, commending at about day 10 of adulthood (**Fig. 8C**). We also find that 5 day old adult animals display a signature change in aging, namely increased age pigment in the gut of adult worms (**Fig. 8D**). In light of these premature aging phenotypes, we surmise that the locomotory defects described above may also be an indication of premature aging.

## DISCUSSION

Together with a parallel study by Reddy and colleagues (Reddy *et al.* 2017), our study provides the first functional characterization of the large, unusual family of *pals* genes in *C. elegans.* In the context of whole animal transcriptome profiling under different conditions, expression of members of the *pals* gene family has previously been shown to be induced upon various forms of cellular insults, ranging from exposure to intracellular fungal infections, to viral infection and to toxic compound exposure (Cui *et al.* 2007; Bakowski *et al.* 2014; Chen *et al.* 2017). We define here a function for one of the family members, *pals-22* in controlling the silencing of repetitive DNA sequences. Even though the biochemical function of PALS proteins is presently unclear, the upregulation of many *pals* genes under conditions of cellular stress suggest that this gene family may be part of a host defense mechanism that protects animals/cells from specific insults. The *C. elegans*-specific expansion of *pals* genes may relate to their potential function in fending off species-specific stressors and/or encounters with species-specific pathogens. Consistent with the species-specificity of *pals* gene function, it has been noted that distinct wild isolates of *C. elegans* display an above-average accumulation of polymorphisms in most *pals* genes (Volkers *et al.* 2013).

The mutant phenotype of *pals-22* as a transgene silencer, as well the dependence of this phenotype on small RNA production, indicates that PALS-22 may control gene expression via small RNA molecules. A role of PALS-22 in controlling gene expression is also illustrated in a parallel study in which *pals-22* has been found to be required for the proper regulation of a battery of stress- and microspiridial infection-induced genes, including many of the *pals* genes themselves (Reddy *et al.* 2017). While the function of *pals-22* in the RNAi process is not clear, there are numerous examples of mutants in which somatic transgene silencing is induced as a result of an increase in RNAi sensitivity, including *rrf-3, eri-1, lin-35* and others (Simmer *et al.* 2002; Kennedy *et al.* 2004; Kim *et al.* 2005; Wang *et al.* 2005; Lehner *et al.* 2006; Fischer *et al.* 2008; Fischer *et al.* 2011; Fischer *et al.* 2013). Since the expression levels per copy of repetitive tandem arrays are much lower than for endogenous genes, transgene targeting siRNAs appear to be already abundant in wild-type transgenic strains, indicating a background level of transgene silencing in wild-type worms (Mello AND Fire 1995). These transgene-targeted siRNAs may be reduced in a mutant displaying enhanced transgene silencing (such as *pals-22*), perhaps because of a shift in the balance between the loading of transgene siRNAs into a silencing Argonaute (*e.g.* NRDE3) versus an anti-silencing Argonaute (*e.g.* CSR-1) (Shirayama *et al.* 2012; Fischer *et al.* 2013).

The precise biochemical function of any PALS protein remains obscure. The only notable sequence relationship that we could find points to potential biochemical function of the PALS proteins in proteostasis. One of the LGIII clusters of *pals* genes (III: -21.90 cM) contains also a large number of *fbxa* genes, another vastly *C. elegans*-specific expanded gene family (Thomas 2006)(**Fig**̤ *fbxa* genes code for F-box proteins that are involved in protein degradation (Thomas 2006). One of the FBXA proteins in the LGIII cluster, FBXA-138, displays sequence similarities to a number of distinct PALS proteins within and outside the LGIII *pals* cluster, including PALS-22, PALS-23, PALS-32, PALS-25 and PALS-1. Even though PALS proteins are not predicted to contain a canonical F-box, it is conceivable that their distant sequence relationship to F box proteins may suggest a role of PALS proteins in protein degradation. How, in the case of PALS-22, such a function may relate to the control of gene expression via small RNA molecules is not clear.

Although it is difficult to unambiguously distinguish accelerated aging from general sickness, young adult *pals-22* mutants clearly exhibit multiple features of aged animals—impaired mobility, elevated age pigments/lipofuscin, and shortened lifespan. Increased expression of repetitive sequences has been documented in aging *C. elegans, Drosophila*, and humans, and has been suggested to contribute to genomic instability and cell dysfunction (Sedivy *et al.* 2013); the physiological effect of decreases in the expression of repetitive sequences has not been explored. However, we note that other transgene silencing mutants (which are also RNAi hypersensitive), like *rrf-3* and *eri-1*, do not display an aging defect (Zhang *et al.* 2009; Ren *et al.* 2012). *pals-22* may therefore be involved in novel aspects of small RNA-dependent gene silencing that may control the expression of genes involved in animal physiology.

## ACKNOWLEDGEMENTS

We thank Chi Chen for the generation of transgenic strains, the CGC for providing strains (funded by NIH Office of Research Infrastructure Programs, P40 OD010440), Kevin Howe and Gary Williams at WormBase for their help obtaining and analyzing protein domain data, Sangeena Salam for initial life span assays, the Million Mutation Project for providing the *pals-19(gk166606) and pals-25(gk891046)* alleles, Emily Troemel for communicating unpublished results, Dylan Rahe for providing the GFP-tagged *che-1(ot856)* allele, HaoSheng Sun for providing the *lin-4^prom^::yfp* MiniMos strain, and members of the Hobert lab for comments on this manuscript. This work was supported by an R01AG046358 to MD, and a NJ Commission on Cancer Research Postdoctoral Fellowship DHFS16PPC070 to GW and the Howard Hughes Medical Institute (OH). E.L.-D. was funded by an EMBO long-term fellowship.

